# Feeding intervention potentiates the effect of mechanical loading to induce new bone formation in mice

**DOI:** 10.1101/2020.12.27.424485

**Authors:** Hasmik Jasmine Samvelyan, John Cummings Mathers, Timothy Michael Skerry

## Abstract

The benefits of increased human lifespan depend upon duration of healthy, independent living; the healthspan. Bone-wasting disorders contribute significantly to loss of independence, frailty and morbidity in older people. Therefore, there is an unmet need globally for lifestyle interventions to reduce the likelihood of bone fractures with age. Although many mechanisms are involved in disorders of bone loss, there is no single regulatory pathway and, therefore, there is no single treatment available to prevent their occurrence. Our aim in these studies was to determine whether fasting/feeding interventions alter the effect of mechanical loading on bone anabolic activities and increase bone mass. In young 17-week-old mice, 16-hour fasting period followed by reintroduction of food for 2 hours increased markedly the potency of mechanical loading, that mimics the effect of exercise, to induce new cortical bone formation. Consistent with this finding, fasting and re-feeding increased the response of bone to a loading stimulus that, alone, does not stimulate new bone formation in ad-lib fed mice. Older mice (20-months) experienced no potentiation of loading-induced bone formation with the same timing of feeding interventions. Interestingly, the pre-, prandial and postprandial endocrine responses in older mice were different from those in young animals. The hormones that change in response to timing of feeding have osteogenic effects that interact with loading-mediated effects. Our findings indicate associations between timing of food ingestion and bone adaptation to loading. If translated to humans, such non-pharmacological lifestyle interventions may benefit skeletal health of humans throughout life-course and in older age.

## Introduction

The dramatic increase in human life expectancy in the last 60 years has been driven by reduced deaths of people in their 60s and 70s, leading to demographic shifts so that most populations globally are characterised by increasing numbers of older old people^1^. However, an increasing number of these extra years of life are spent in poor health with more years spent in care facilities due to reduced ability to live independently. Osteoporosis is a major contributor to loss of independence due to bone fractures, and resulting hospital treatments lead to significant morbidity. Even after successful fracture treatment, independent living is compromised in many patients^2^. While drug treatments reduce consequences of osteoporosis significantly, there is a pressing need for non-pharmacological interventions to improve bone health across the life-course and to reduce likelihood of age-related bone disease.

Physical activity is beneficial for the skeleton acting through the process of functional adaptation, a process that is less effective in older animals and humans than young ones^3–6^. In addition, older people and those with bone loss, who are already at increased risk of fractures, may be unable to perform exercise that is sufficiently vigorous to strengthen their bones. Since dietary interventions such as energy (caloric) restriction are associated with healthy ageing, a combination of exercise and nutrition may benefit bone health in ageing by utilising synergies between osteotropic influences.

Increased physical activity is beneficial for the skeleton by reducing the rate of age-related loss of bone mass through shifting the balance of bone remodelling in favour of bone formation^7^. However, during ageing skeletal resorption rates are increased with decrease in bone formation rates, resulting in overall net bone loss^8^. Therefore, it is hard for elderly adults and patients with bone wasting disorders to perform vigorous exercise safely, and there is a need to identify effective ways for those people to exercise which produce maximum benefits for the musculoskeletal system without increased risk of injury.

There is complex and coordinated relationship between the endocrine regulation of energy metabolism, adipose tissue and bone homeostasis^9–12^. Specifically, concentrations of hormones including ghrelin, leptin, insulin, amylin, incretins, GIP, GLP1 and GLP2 change profoundly in anticipation of, during or after eating (preprandial, prandial and postprandial responses). These hormones have anabolic effects on bone homeostasis, therefore, synergies between endocrine and mechanical effects could alter with changes in concentrations of these hormones. Indeed, previous studies in model organisms provide evidence of synergistic effects of one of the osteoregulatory hormones; parathyroid hormone and mechanical loading on bone. Parathyroid hormone administration alone induces bone formation but, in combination with mechanical loading, enhances mechanically-induced bone formation synergistically^13–17^.

In this study, we have investigated the synergistic effect of feeding interventions following an overnight fast on the response of bone to mechanical loading in mice and the concurrent role of candidate osteotropic hormones in bone homeostasis. Using a well-established tibial axial loading model^18,19^, we performed experiments to determine osteogenic effects of different mechanical loading regimens after periods of withholding food (fasting) or feeding in young (17-19 week-old) and aged (20-month-old) male C57BL/6 mice. We measured new bone formation in these mice using micro-computed tomography and dynamic histomorphometry. To explore possible mechanisms by which fasting/feeding interventions could have their effect, we measured changes in serum concentrations of a range of candidate osteotropic hormones in respone to feeding in fed and fasted fed young and aged mice using a multiplex assay.

## Materials and Methods

### Animals

Male C57BL/6 wild type mice at 16 weeks of age (young adult) and 20 months of age (aged) were obtained from Charles River Laboratories Inc. (Margate, UK). The mice were caged separately and acclimatised to their surroundings for 7 days. All mice were allowed free access to water and a maintenance diet ad libitum (2018C Teklad Global 18% Protein Rodent Diet; Madison, WI, USA, Supplementary Table S1) in a 12-hour light/dark cycle at a room temperature of 21 ± 2°C and relative humidity of 55 ± 10%. Cages contained wood shavings, bedding and metal rings for mice to play. All procedures complied with the United Kingdom Animals (Scientific Procedures) Act 1986 (ASPA) and were reviewed and approved by the University of Sheffield Research Ethics Committee (Sheffield, UK). 3Rs principle followed as the ethical framework for conducting all the scientific experiments.

### Assessment of feeding behaviour of fed or fasted young and aged mice

Groups of young adult male mice at 17 weeks of age and aged male mice at 20 months of age were either allowed access to a maintenance diet ad libitum or fasted overnight for 16 hours and were assessed for subsequent feeding behaviour. The nesting material, bedding and wood shavings were removed from the cages prior to the experiment. All mice were allowed free access to water. Food intake was measured by monitoring the consumed food and body weights of animals every 30 minutes for the first 3 hours then every hour for another 4 hours twice weekly for one week. After the experiments mice were placed in new cages with wood shavings and bedding.

### In vivo non-invasive axial mechanical loading of knee joints

The right tibia of each mouse was subjected to non-invasive, dynamic axial mechanical loading under the isoflurane-induced anaesthesia (liquid isoflurane was vaporised to a concentration of 5% and maintained at a concentration of 3% with oxygen) for 7min/day, 3 alternate days a week for 2 weeks according to the protocols described in the previous studies^16,18^. The left tibiae were non-loaded internal controls. To apply mechanical loading, mice were anaesthetised, and the knee and ankle joints of the right limbs were fixed in customised concave cups. The knee was positioned into the upper cup, which was attached to the activator arm displacement transducer and the ankle in the lower cup attached to the dynamic load cell. The tibia was held in place by continuous static preload of 0.5N onto which dynamic loads were superimposed in a series of 40 trapezoidal shaped waveform cycles with steep up and down ramps with a high dwell time of 0.25s and 9s low dwell ‘rest interval’ between each cycle. The load was applied to engender the required magnitude of strain on the bones of animals in each of the age groups^20^. The engendered average strain rate was 30000με/s, which is shown to be equivalent to physiological strain rates^21^.

### Calcein administration

150mg of calcein powder was added to 50ml dH_2_O. Then 0.1g of NaHCO_3_ was added to neutralise the pH of the solution. The solution was protected from light with aluminium foil. After the solution was fully dissolved, it was filter-sterilized at 0.22μm into autoclaved vials also protected from light with aluminium foil. The pH of the solution was measured with a bench pH-meter and corrected to neutral pH 7 with HCl or NaOH. The calcein solutions were made on the same days prior to the experiments ensuring no loss of fluorescence from the fluorochrome label. Calcein was administered at a concentration of 30mg/kg in 0.2% NaHCO_3_ solution subcutaneously on the first and last days of mechanical loading (days 1 and 12) using insulin syringes^22^.

### Micro-computed tomography (μCT) analysis

The tibiae were collected after mice were culled, and loading responses of bones determined by μCT using a SkyScan 1172 (Bruker MicroCT, Kontich, Belgium). Since mouse bones are small, and axial loading related osteogenesis is site-specific, high-resolution μCT analysis was used primarily to quantify three-dimensional (3D) bone microarchitecture at comparable sites (proximal and mid-shaft regions) of loaded and contralateral non-loaded control tibiae. The tibiae were stored in 70% EOH and mounted in a plastic tube wrapped in cling film to prevent drying during scanning. The high-resolution scans were imaged with a pixel size of 4.3μm. The applied X-ray voltage was 50kV, X-ray intensity 200μA with a 0.5mm aluminium filtration. The scans were taken over 180 degrees with a 0.7-degree rotation step. The images were reconstructed and binarised with a threshold of 0 to 0.16, ring artefact reduction was set at 10 and beam hardening correction at 0% using the SkyScan NRecon software package (version 1.6.9.4, Bruker MicroCT, Kontich, Belgium). The images then were realigned vertically using DataViewer software (version 1.5.1.2 64-bit) before the 3D quantification.

### 3-dimensional (3D) and colour coded analysis

Structural parameters were calculated using SkyScan Ct.An software (Bruker MicroCT, Kontich, Belgium) for cortical bone 1-mm-long section at the mid-shaft of the tibia with an offset of 0.5mm, and for trabecular bone - secondary spongiosa, 1mm distal to the tibial proximal growth plate with an offset of 0.2mm ensuring the growth plate was not included in the analysis. In the trabecular region, an irregular, anatomic region of interest (ROI) adjacent to the endocortical boundary was analysed. Two ROIs were drawn to analyse the cortical region including ‘doughnut’ shaped ROI and ‘crude’ ROI around the outside of the bone.

The bone mass, architecture and changes due to the mechanical loading were evaluated in cortical and trabecular bone regions according to the guidelines for the assessment of bone microstructure in rodents using μCT^23^. 3D representations of the bones were created using the Voxler software (Golden Software Inc., 2006). The local bone thickness (Th; voxel) was determined and colour-coded thickness images were generated using the Avizo® (version 8.0, VSG, Burlington, VT, USA) software.

### Bone dynamic histomorphometry

After μCT scanning and being fixed in 70% EOH for a minimum of 48hrs, the mouse bones were infiltrated in increasing concentrations of alcohol (80% IMS alcohol, 80% IMS alcohol, 100% IMS alcohol, 100% IMS alcohol, 100% EOH, 100% EOH and 100% EOH) on a shaker at room temperature for 48hrs at each concentration. Bones were then infiltrated in glass bottles with medium grade LR White resin at 4°C for 6 days, resin was changed every 48hrs. The sample bottles were agitated gently and continuously on a shaker for a duration of the infiltration at 4°C. Next, the bones were placed longitudinally and diagonally in plastic moulds fully filled with fresh resin and covered with labelled plastic embedding stubs at room temperature. Resin was then allowed to polymerise at 55°C overnight for up to 24hrs until it was hard. After polymerisation, blocks were removed from the moulds and protected from the light during the storage and subsequent use. The bones were protected from the light at all stages of the processing to prevent the loss of fluorescence from the labels.

### Sectioning and visualisation

The bones were reoriented vertically, then 8μm thick transverse sections were obtained from the cortical ROI corresponding to the mid-shaft of the tibiae and as determined by μCT, using the tungsten carbide-tipped 16cm microtome knife on the rotary microtome. Sections were unrolled and floated onto a drop of the dH_2_O on a labelled Superfrost Plus glass slides using fine paintbrush and forceps. The slides were covered with Saran wrap and wrapped with blotting paper making sure the Saran wrap-covered sections were flat without creases or residual water. The slides then were dried at 50°C overnight stacked in the slide clump to aid adherence of bones sections. After peeling off the Saran wraps slides were mounted in the xylene for 2-3min and then covered with glass coverslip using DPX mountant. Images of calcien labelled transverse bone sections were visualised using Leica fluorescent microscope at 4x and 10x magnifications. All slides were stored protected from the light until time for histomorphometry analysis.

### Measurements of bone formation parameters

The histological assessment of bone phenotypes was carried out by dynamic histomorphometry measuring its derived kinetic indices^24^. A Leica microscope was set up for fluorescence before placing the slides with calcein labeled sections under the microscope. To reach its full output, the fluorescent lump was allowed to warm up for 5min before measurements were made. Bone formation parameters including mineralising surface (MS), mineral apposition rate (MAR) and bone formation rate (BFR) were measured using the OsteoMeasure software (version 3.3.0.2).

### Blood collection

Small blood volumes were obtained from the mice by tail vein sampling technique with assistance from the Biological Services Unit at the Royal Hallamshire Hospital of The University of Sheffield. To avoid hyperthermia and dehydration and reduce stress levels, this technique was performed without warming mice in the warming cabinet prior to taking the blood samples. Mice were gently placed in the restraint tube and lateral tail veins were accessed by making incisions in approximately one-third from the proximal end along the length of the tails^25^. Repeated blood micro samples (typically ≤50μl) were collected from the lateral mouse-tail veins 3 times in any one 24-hour period.

### Mouse multiplex metabolic hormone assay

Serum ghrelin, leptin, insulin, GIP and GLP1 concentrations were assayed in mouse sera using mouse multiplex assay (Bio-Rad, Watford, UK) following manufacturer’s instructions. The mouse assay is magnetic bead-based multiplex assay that allows measurement of multiple proteins in small volumes of sera. Briefly, a 96-well plate was pre-coated with 50μl of beads coupled with the capture antibodies. After washing twice with wash buffer, 50μl of standards, blank and serum samples were added into the wells, incubated at room temperature with shaking for 1hr, washed and followed by addition of the detection antibodies and streptavidin-phycoerythrin (SA-PE) conjugates. Following incubation at room temperature with shaking, the plate was washed three times with wash buffer and beads were resuspended in 125μl assay buffer with shaking at 850rpm for 30s. The plate was then read on Bio-Plex system using Bio-Plex Manager software (v 6.0). Samples (pg/ml) were assessed in duplicate.

### Serum cortisol analysis

Cortisol concentrations were measured in mouse sera by a solid phase competitive enzyme linked immunosorbent assay (ELISA) (Sigma-Aldrich Co Ltd, USA) according to the manufacturer’s protocol. Prior to the assay the frozen sera were thawed at a room temperature and mixed well using vortexer. Briefly, 25μl of mouse serum was added into each microwell of the plate, coated with the anti-cortisol MAb, along with the standards for 6-point standard curve (1-80ng/ml), blank and internal control. After incubation at room temperature with shaking for 60min, wells were washed three times with wash buffer and treated with 3,3’,5,5’ - Tetramethylbenzidine (TMB) substrate for 15min. The colorimetric reaction was then stopped and absorbance was measured at 450nm on ELISA reader. A 4-parametre logistic regression was applied to the standard curve for the calculation of cortisol concentrations in the samples (ng/ml). Samples were assessed in duplicate.

### Statistical analysis

All analyses were performed using GraphPad Prism software 6.0f version (GraphPad Inc, La Jolla, CA, USA). The results were presented as the mean ± standard error (SEM). The Normal distribution of data was assessed using the Shapiro-Wilk normality test. Statistical significance was tested using parametric statistical tests. For comparing two groups, two-tail Student’s *t*-test (paired or unpaired) was used. For comparing more than two groups, two-way ANOVA (analysis of variance) was used with Tukey post-hoc test to assess the effect of loading, fasting/feeding and interactions within and across the experimental groups. The significance was set at *p* < 0.05.

## Results

### The effect of 16-hour food restriction on feeding behaviour of male mice

To establish a robust model to study the effect of timing of feeding on bone’s response to loading, we first assessed the effect of overnight, 16-hour food removal on subsequent feeding behaviour of young (17-18 weeks old) and aged (20-month-old) male C57BL/6 mice. The results provided insights into dynamics of the homeostatic feeding response and showed that both young adult (Figure 1a) and aged (Figure 1b) male mice exhibited a similar 2-hour hyperphagic response following fasting before returning to the same food intake as mice that had unrestricted access to a normal chow ad libitum for the whole period of experiment (Figure 1a, b).

**Figure 1.**
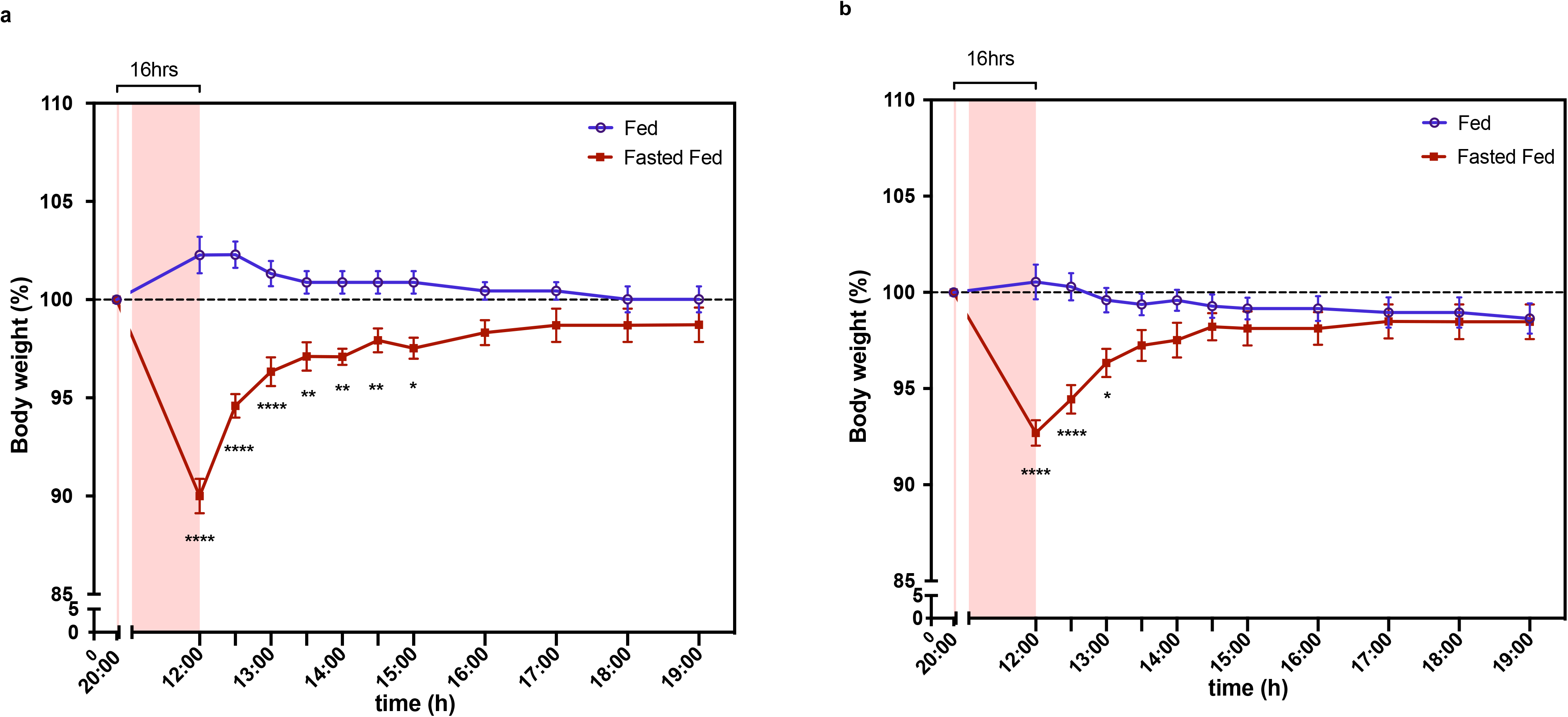
Feeding behaviour of young and aged male C57BL/6 mice. **a** In total 14 mice were randomly divided into two groups; control ad-lib fed (*n* = 7) or overnight 16-hour fasted (*n* = 7) at 17 weeks of age. **b** In total 16 male mice were randomly divided into two groups; ad-lib fed (*n* = 8) or 16-hour fasted (*n* = 8) at 20 months of age. The body weights of mice were measured for 7 hrs following overnight fast to monitor food intake of the animals. Following overnight food restriction, fasted mice consumed more food compared with the mice in the control group that had free access to food ad libitum. The body weights of both young adult and aged mice in the fasted group increased significantly for 2 hrs compared with their pre-fasted body weights indicating rapid ingestion of food (and water) in the early re-feeding period. The proportions of the starting pre-fasted body weights (%) ± SEM are shown. * indicates significant differences between fed and fasted fed groups; * *p* < 0.05, ** *p* < 0.01, **** *p* < 0.0001 by two-way ANOVA with Tukey post-hoc test

### Feeding intervention potentiates adaptive response of the bones to mechanical loading in young mice

Next we compared bone formation responses in ad-lib fed animals and in mice fasted overnight and then allowed free access to maintenance diet for 1, 2 or 3 hours before loading. To mimic osteogenic exercise, compressive forces were applied between the hock (ankle) and stifle (knee) joints to induce deformation of the tibiae. Different loading regimens comprised 40 cycles of compressive force at high physiological strain rate to peak strain magnitudes of 2200, 1300 or 1100 microstrain, followed by a brief hold of 0.25 seconds, and a rapid symmetrical reduction in loading, with a 9 second rest period before the next compression. The three strain magnitudes corresponded to maximal, sub-maximal and sub-threshold stimuli respectively. Loading was applied on Monday, Wednesday and Friday of 2 successive weeks, at 1, 2 or 3 hours after food had been re-introduced to 16-hour fasted mice, or at the same time of day in ad-lib fed animals (Supplementary Figure S1).

Using a high resolution μCT analysis of the cortical bone at the mid-shaft region of loaded tibiae, we found the expected responses to loading at high physiological strains with significant adaptive response in the mice fed ad-lib. However, in mice that were fasted then fed for 2 hours, the change in cortical bone thickness induced by maximal loading was increased by 36% compared with responses of mice fed ad-lib before the loading (0.049 ± 0.007mm in fasted/fed loaded tibiae compared with that in ad-lib fed mice 0.036 ± 0.004mm, *p* < 0.05) (Figure 2a, c, d). The histomorphometric data were consistent with the μCT measurements (Figure 2b, Table 1). Colour-coded analysis revealed anatomical variation in cortical bone thickness between loaded and non-loaded tibiae of fasted fed and ad-lib fed mice (Figure 2d). Differences in the cortical bone thickness of loaded tibiae were particularly apparent between fasted/fed and ad-lib fed mice and correlated with μCT analysis. There was no significant induction of bone formation in non-loaded legs of either ad-lib or fasted/fed mice over the period of the experiment.

**Figure 2.**
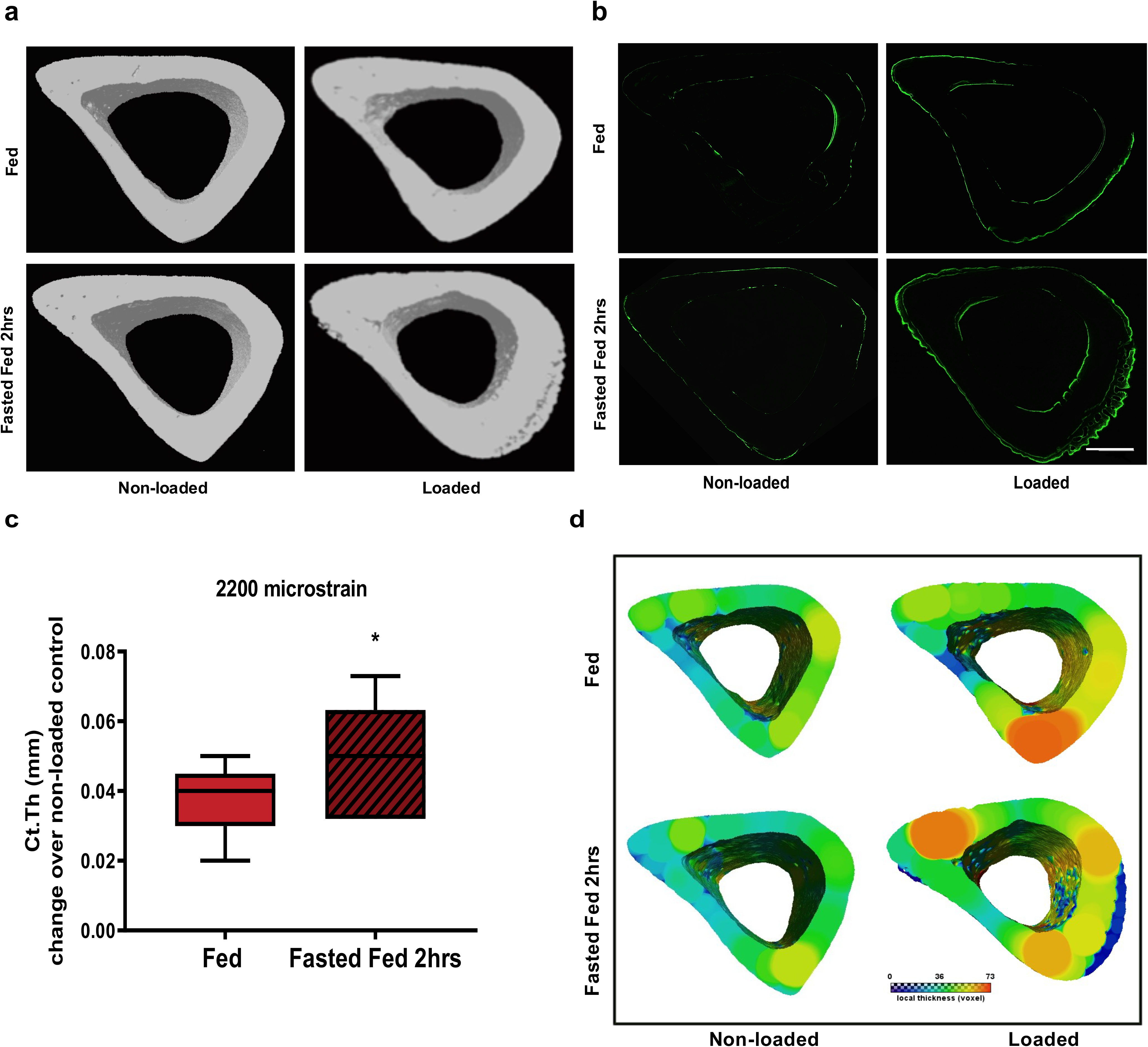
Cortical bone phenotype of control (non-loaded) and loaded tibiae with 2200 microstrain (maximal) loading regimen in 19-week-old fed or overnight fasted 2-hour fed male C57BL/6 mice. **a** Three-dimensional images of a 1.0-mm-thick cortical bone region 0.9mm below the growth plate from the mid-shaft of control non-loaded (left) and loaded (right) tibiae of fed or fasted 2-hour fed mice. **b** Transverse cortical bone confocal microscope images from the mid-shaft of left non-loaded and right loaded tibiae of fed (top) or fed for 2 hours following overnight fast (bottom) male mice. Double calcein labels can be visualised on endocortical and periosteal surfaces of the loaded tibiae confirming new bone formation at these sites. **c** The change in cortical bone thickness (Ct.Th) of tibiae loaded at 13N to induce peak strain magnitude of 2200 microstrain compared with non-loaded controls of fed or fasted 2-hour fed young adult male mice measured using μCT at tibial mid-shaft. Fasted 2-hour fed mice had significantly greater response in Ct.Th of the right tibiae compared with ad-libitum fed mice. Scale bar = 200μm. **d** Three-dimensional visualisation and representative colour-coded images of the cortical bone thickness of non-loaded and loaded tibiae of fed and fasted 2-hour fed young adult mice. All data are means ± SEM. *n* = 7 fed mice, *n* = 6 fasted 2-hour fed mice. **c** Two-tailed Student’s *t*-test; * *p* < 0.05

**Table 1.**
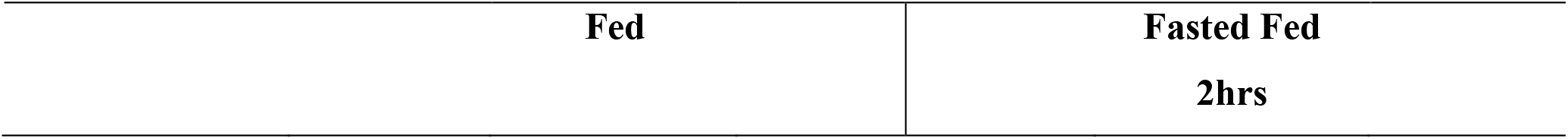
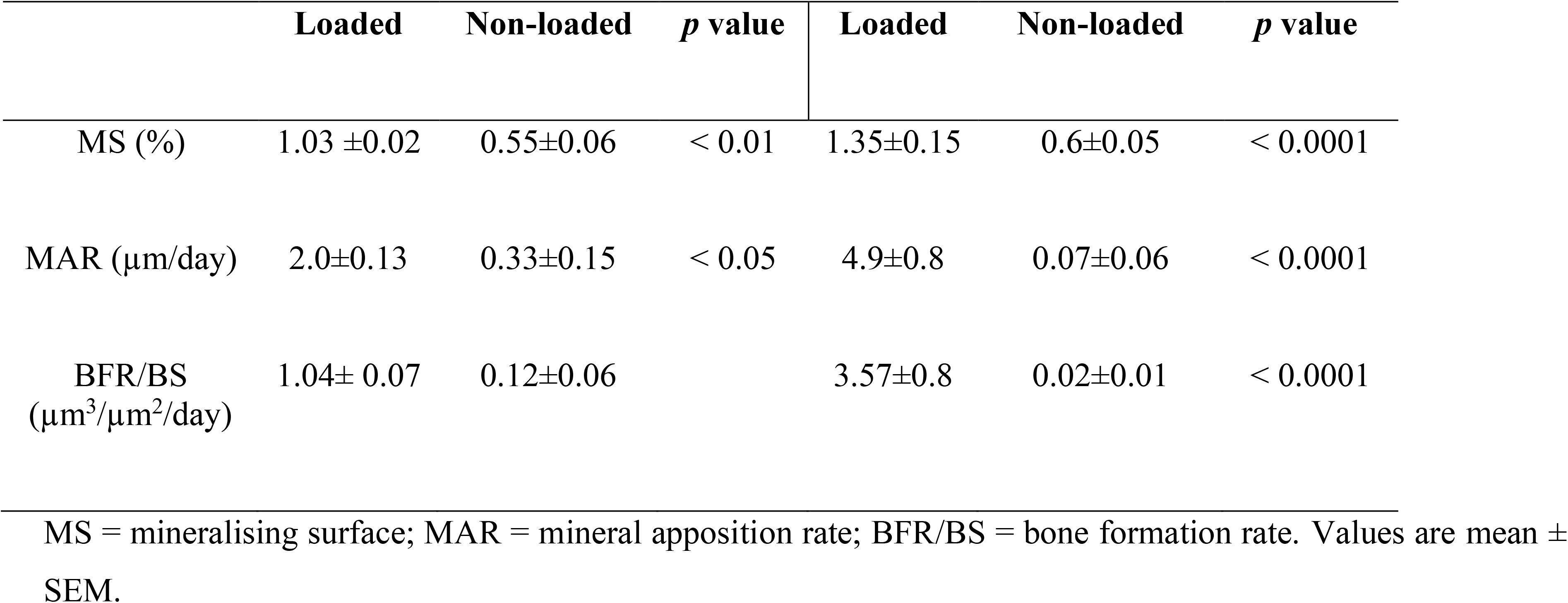
Effect of 2-hour feeding following overnight fast on load-induced cortical bone formation in young adult mice.

|  | Loaded | Non-loaded | *p* value | Loaded | Non-loaded | *p* value |
| --- | --- | --- | --- | --- | --- | --- |
| MS (%) | 1.03 ±0.02 | 0.55±0.06 | < 0.01 | 1.35±0.15 | 0.6±0.05 | < 0.0001 |
| MAR (µm/day) | 2.0±0.13 | 0.33±0.15 | < 0.05 | 4.9±0.8 | 0.07±0.06 | < 0.0001 |
| BFR/BS<br>(µm<sup>3</sup>/µm<sup>2</sup>/day) | 1.04± 0.07 | 0.12±0.06 |  | 3.57±0.8 | 0.02±0.01 | < 0.0001 |

MS = mineralising surface; MAR = mineral apposition rate; BFR/BS = bone formation rate. Values are mean ± SEM.

In animals fed for 1 or 3 hours after the end of fasting, the mean cortical thickenss increased compared with ad lib-fed controls, but the difference was not significant. Trabecular bone parameters were improved by loading in both fasted and ad-lib fed groups and at all 3 times after re-introduction of food, but were not significantly different between feeding interventions.

### Load magnitude-related adaptive response of the cortical bone is potentiated by feeding intervention in young mice

We then determined effects of fasting and 2-hour feeding regimen on bone response to loading which, in ad-lib fed mice, induced less than maximal bone formation. In mice, whose bones were loaded to induce peak strain magnitudes of 1300 microstrain, a sub-maximal stimulus, loading after fasting and feeding increased cortical thickness by 13% compared with ad-lib fed mice (fasted/fed: 0.27 ± 0.007mm loaded limb versus 0.24 ± 0.003mm control limb, *p* < 0.05; ad-lib fed: 0.26 ± 0.004mm versus 0.24±0.004mm, *p* < 0.05) (Figure 3b). Finally, we investigated the effects of a sub-threshold stimulus (1100 microstrain), not capable of inducing new bone formation in ad-lib fed mice. Remarkably, that previously ineffective stimulus together with imposition of the fasting/re-feeding regimen resulted in an 8% increase in cortical thickness (fasted/fed: 0.26 ± 0.008mm loaded limb versus 0.24 ± 0.008mm control limb, *p* < 0.05; ad-lib fed: 0.26 ± 0.002mm versus 0.25 ± 0.004mm, *p* > 0.05) (Figure 3a). We did not detect significant changes in trabecular bone parameters in these studies.

**Figure 3.**
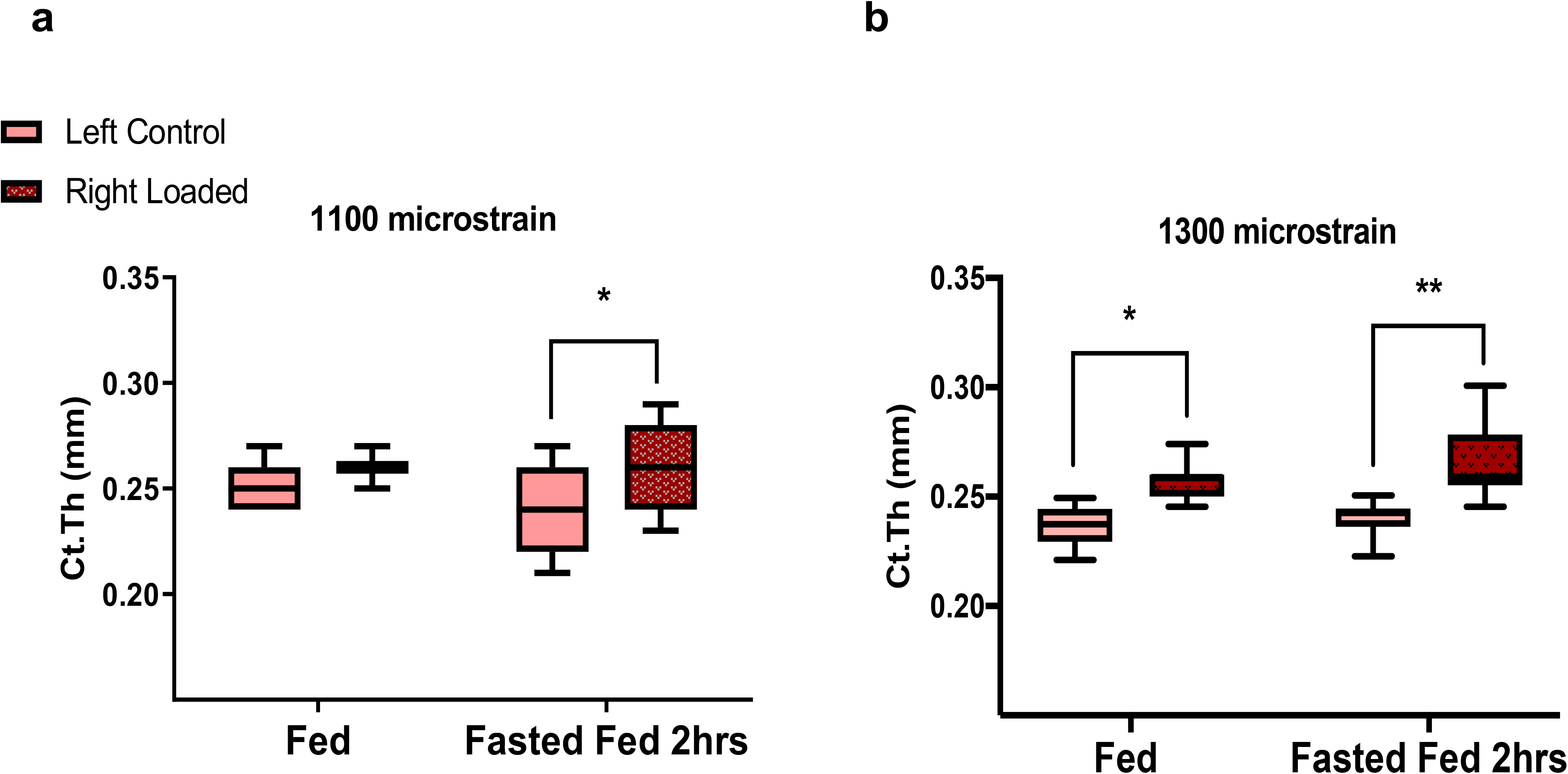
Cortical bone phenotype of control (non-loaded) and loaded tibiae with 1100 microstrain (sub-threshold) and 1300 microstrain (submaximal) loading regimens in 19-week-old fed or overnight fasted 2-hour fed male C57BL/6 mice. **a** The effect of sub-threshold and **b** submaximal mechanical loadings with peak force of 8N to induce peak strain magnitude of 1100 microstrain and 11N to induce peak strain magnitude of 1300 microstrain respectively on Ct.Th of fed or fasted 2-hour fed young adult mice. All data are means ± SEM, *n* = 7. **a, b** Two-way ANOVA with post-hoc Tukey comparisons; * *p* < 0.05, ** *p* < 0.01

### Adaptive response of bones to mechanical loading in aged mice is not potentated by the same feeding intervention that is effective in young mice

We then compared effects of fasting/feeding regimens on load-induced bone formation in older 20-month-old mice, but observed no potentiation effect of loading of the fasting and 2 hour feeding regimen that we had seen in younger mice (cortical bone thickness fasted/fed: 0.23 ± 0.002mm loaded limb versus 0.21 ± 0.004mm control limb, *p* < 0.05; ad-lib fed: 0.24 ± 0.01mm versus 0.21 ± 0.007mm, *p* < 0.05) (Figure 4a-c). Colour-coded analysis revealed no anatomical variation in cortical bone thickness between loaded and non-loaded tibiae of fasted fed and ad-lib fed aged mice (Figure 4c).

**Figure 4.**
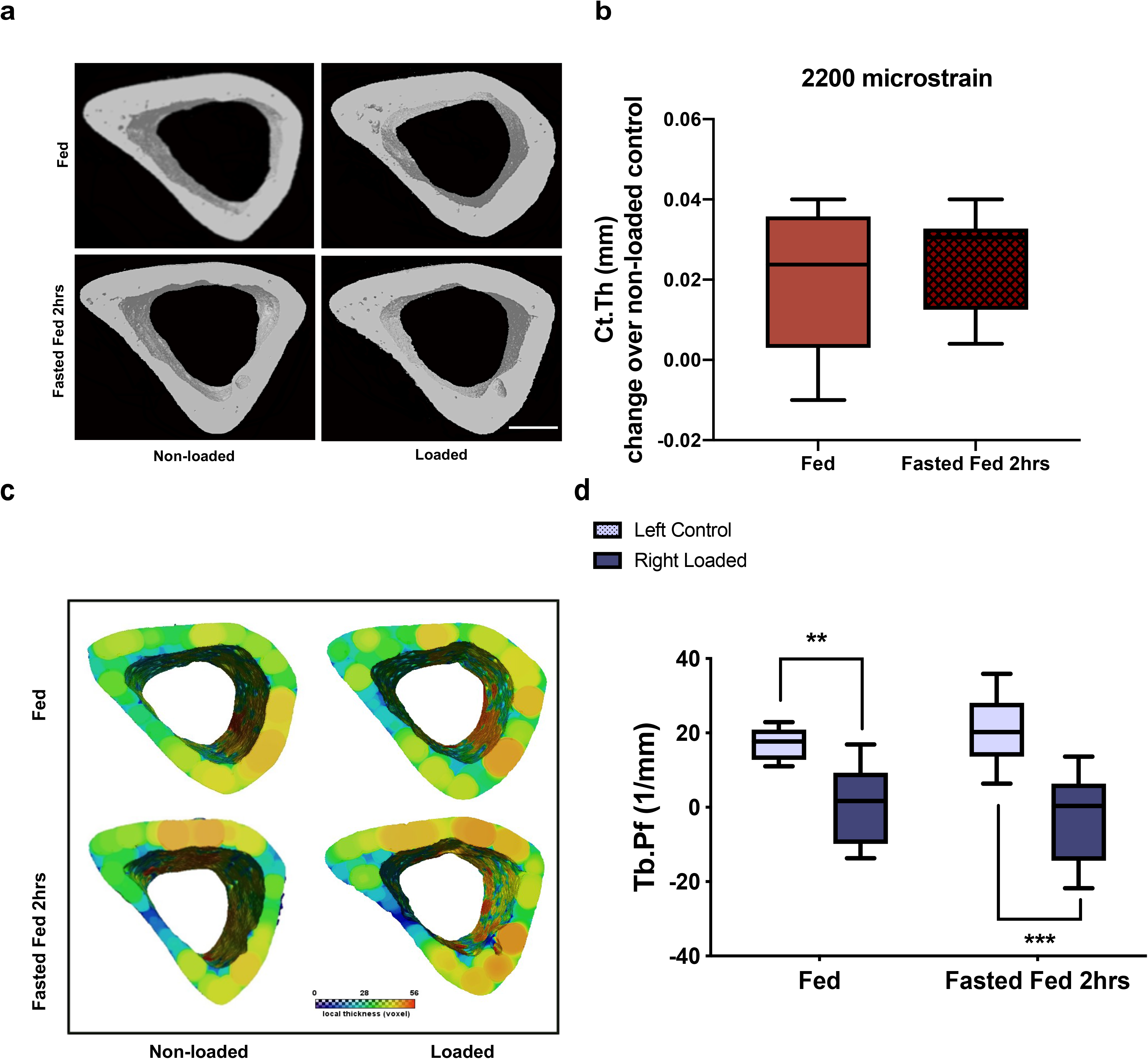
Cortical and trabecular bone phenotypes of control (non-loaded) and loaded tibiae with 2200 microstrain osteogenic loading regimen in 20-month-old fed or overnight fasted 2-hour fed male C57BL/6 mice. **a** Three-dimensional images of a 1.0-mm-thick cortical bone region 0.9mm below the growth plate from the mid-shaft of control non-loaded (left) and loaded (right) tibiae of fed or fasted 2-hour fed aged mice; scale bar = 200μm. **b** The change in cortical bone thickness (Ct.Th) of tibiae loaded at 10N to induce peak strain magnitude of 2200 microstrain compared with non-loaded controls of fed or fasted 2-hour fed aged male mice measured using μCT at tibial mid-shaft. There was no significant difference in Ct.Th of the loaded tibiae of fasted 2-hour fed mice compared with ad-libitum fed mice. **c** Three-dimensional visualisation and representative colour-coded images of the cortical bone thickness of non-loaded and loaded tibiae of fed and fasted 2-hour fed aged adult mice. **d** Trabecular bone pattern factor (Tb.Pf) of the right tibiae loaded at 10N to induce peak strain magnitude of 2200 microstrain compared with non-loaded left controls of fed or fasted 2-hour fed aged male mice measured at tibial proximal region using μCT. Box-plots represent means ± SEM. *n* = 8. **b** Two-tailed Student’s *t*-test **d** Two-way ANOVA with post-hoc Tukey comparisons; ** *p* < 0.01, *** *p* < 0.001

Despite deterioration of the trabecular bone in these aged mice, trabecular bone pattern factor decreased significantly in both fed and fasted 2-hour fed mice (fasted/fed: −2.3 ± 4.3mm^−1^ loaded limb versus 20.8 ± 3.3mm^−1^ control limb, *p* < 0.001; ad-lib fed: 0.7 ± 3.7mm^−1^ versus 17.3±1.5mm^−1^, *p* < 0.01) (Figure 4d). Similarly, there was no significant change of bone formation in non-loaded legs of either ad-lib or fasted/fed aged mice over the period of the experiment.

### Regulation of bone homeostasis by endocrine hormones differ between young and aged mice

Next we assessed changes in circulating concentrations of gastro-entero-pancreatic hormones in response to fasting and feeding, which may contribute to observed potentiating effects. We measured concentrations of a range of hormones with known effects on bone homeostasis in young and old mice, in both ad-lib fed and fasted/fed groups. Since both young and aged mice previously exhibited a similar 2-hour hyperphagic response following fasting before returning to the same food intake, serum concentrations of ghrelin, leptin, pancreatic hormone insulin, and intestinal peptides GIP and GLP1 were measured at 3 time points (overnight 16-hour fasted at 12:00 noon on the day following the fast, then 2 hours and 3 hours after initiating food intake at 14:00 and 15:00 respectively) using a multiplex metabolic assay. Endocrine profiles differed between young and aged mice. Fasting and feeding changed circulating serum concentrations of ghrelin, leptin, insulin and GLP-1, but not GIP in young mice (Table 2). In older mice, absolute concentrations of hormones were different, but differences between ad-lib fed and fasted fed mice were not significant (Table 3).

**Table 2.** Serum concentrations of hormones altered during fasting and feeding in young mice.

|  | Fed |  |  | *p* value |
| --- | --- | --- | --- | --- |
|  | 12:00 | 14:00 | 15:00 |  |
| **Hormone** |  |  |  |  |
| Ghrelin | 703±162 | 1493±156 | 1326±119 | < 0.05<sup>a</sup> |
| Leptin | 298±0.1 | 269±25 | 276±25 |  |
| Insulin | 929±94 | 1023±133 | 1291±110 |  |
| GIP | 54±3 | 50±3 | 51±2 |  |
| GLP1 | 8±1 | 7±1 | 7±1 |  |
|  | **Fasted Fed** |  |  |  |
| Ghrelin | 1233±251 | 757±95<sup>1</sup> | 1143±48 |  |
| Leptin | 95±15<sup>1</sup> | 437±62 | 601±76<sup>3</sup> | < 0.0001<sup>b</sup> |
| Insulin | 552±0.1 | 2175±303<sup>2</sup> | 1319±107 | < 0.05<sup>c</sup> |
| GIP | 44±2 | 58±2 | 56±3 | 0.01<sup>b</sup> |
| GLP1 | 7±1 | 10±1 | 11±1.<sup>1</sup> | 0.05<sup>b</sup> |

Two-way ANOVA used to assess the differences within and between the groups. Comparisons of time points within the same group are indicated as <sup>a</sup>12:00 < 14:00; <sup>b</sup>12:00 < 14:00,15:00; <sup>c</sup>12:00 < 14:00 > 15:00. Comparisons of same time points between the groups are indicated as <sup>1</sup>*p* < 0.05; <sup>2</sup>*p* < 0.01; <sup>3</sup>*p* < 0.001. Samples were assessed in duplicate. Results are expressed as pg/ml, mean ± SEM.

**Table 3.**
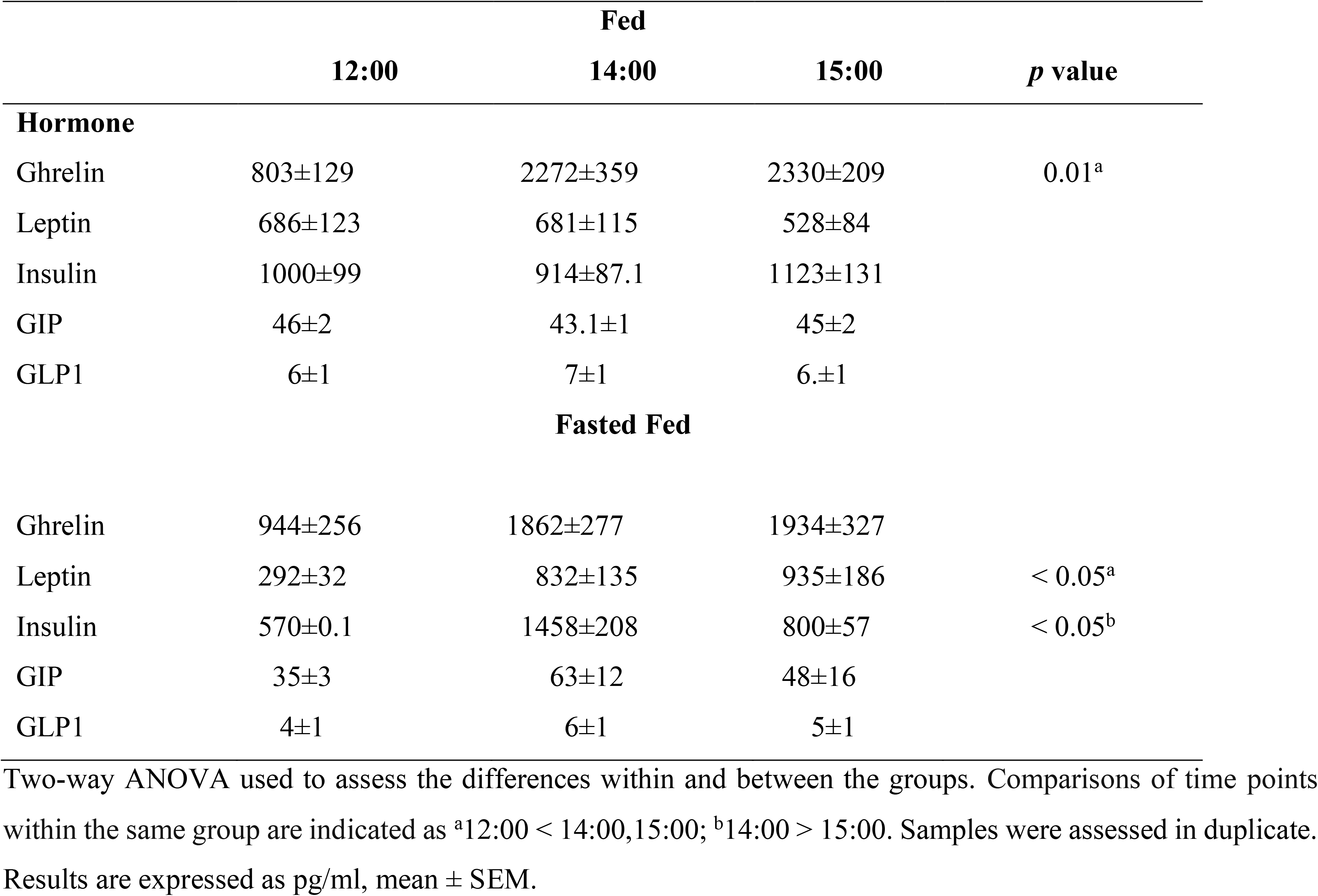
Serum concentrations of hormones altered during fasting and feeding in aged mice.

|  | Fed |  |  | *p* value |
| --- | --- | --- | --- | --- |
|  | 12:00 | 14:00 | 15:00 |  |
| **Hormone** |  |  |  |  |
| Ghrelin | 803±129 | 2272±359 | 2330±209 | 0.01<sup>a</sup> |
| Leptin | 686±123 | 681±115 | 528±84 |  |
| Insulin | 1000±99 | 914±87.1 | 1123±131 |  |
| GIP | 46±2 | 43.1±1 | 45±2 |  |
| GLP1 | 6±1 | 7±1 | 6±1 |  |
|  | **Fasted Fed** |  |  |  |
| Ghrelin | 944±256 | 1862±277 | 1934±327 |  |
| Leptin | 292±32 | 832±135 | 935±186 | < 0.05<sup>a</sup> |
| Insulin | 570±0.1 | 1458±208 | 800±57 | < 0.05<sup>b</sup> |
| GIP | 35±3 | 63±12 | 48±16 |  |
| GLP1 | 4±1 | 6±1 | 5±1 |  |

Two-way ANOVA used to assess the differences within and between the groups. Comparisons of time points within the same group are indicated as <sup>a</sup>12:00 < 14:00,15:00; <sup>b</sup>14:00 > 15:00. Samples were assessed in duplicate. Results are expressed as pg/ml, mean ± SEM.

Ghrelin concentrations fell significantly 2 hours after initiating food intake in young mice, but remained high in older animals (Figure 5a, b). In fasted fed young and aged mice, serum leptin concentrations increased significantly at 14:00 and 15:00 compared with those at 12:00, whereas in ad-lib fed mice remained unchanged. Fasting serum concentration of leptin was significantly lower at 12:00 and significantly higher at 15:00 in fasted fed young mice compared to those in ad-lib fed mice (Figure 5c and d). In fasted fed young mice serum insulin concentration significantly increased from 12:00 to 14:00 and in comparison to that in ad-lib fed mice, then significantly decreased at 15:00 (Figure 5e). In fasted fed aged mice, serum insulin concentration increased but not significantly from 12:00 to 14:00, then significantly decreased at 15:00 compared to that at 14:00, whereas serum insulin concentrations remained unchanged in ad-lib fed aged mice (Figure 5f). In fasted fed young mice, serum GIP and GLP1 concentrations significantly increased at 14:00 and 15:00 compared with those at 12:00 and remained unchanged in ad-lib fed mice (Figure 5g, i). Serum GLP1 concentration was significantly higher at 15:00 in fasted fed mice compared to that in ad-lib fed mice (Figure 5i). There were no significant changes in serum GIP and GLP1 concentrations within and between fed and fasted fed aged mice (Figure 5h, j).

**Figure 5.**
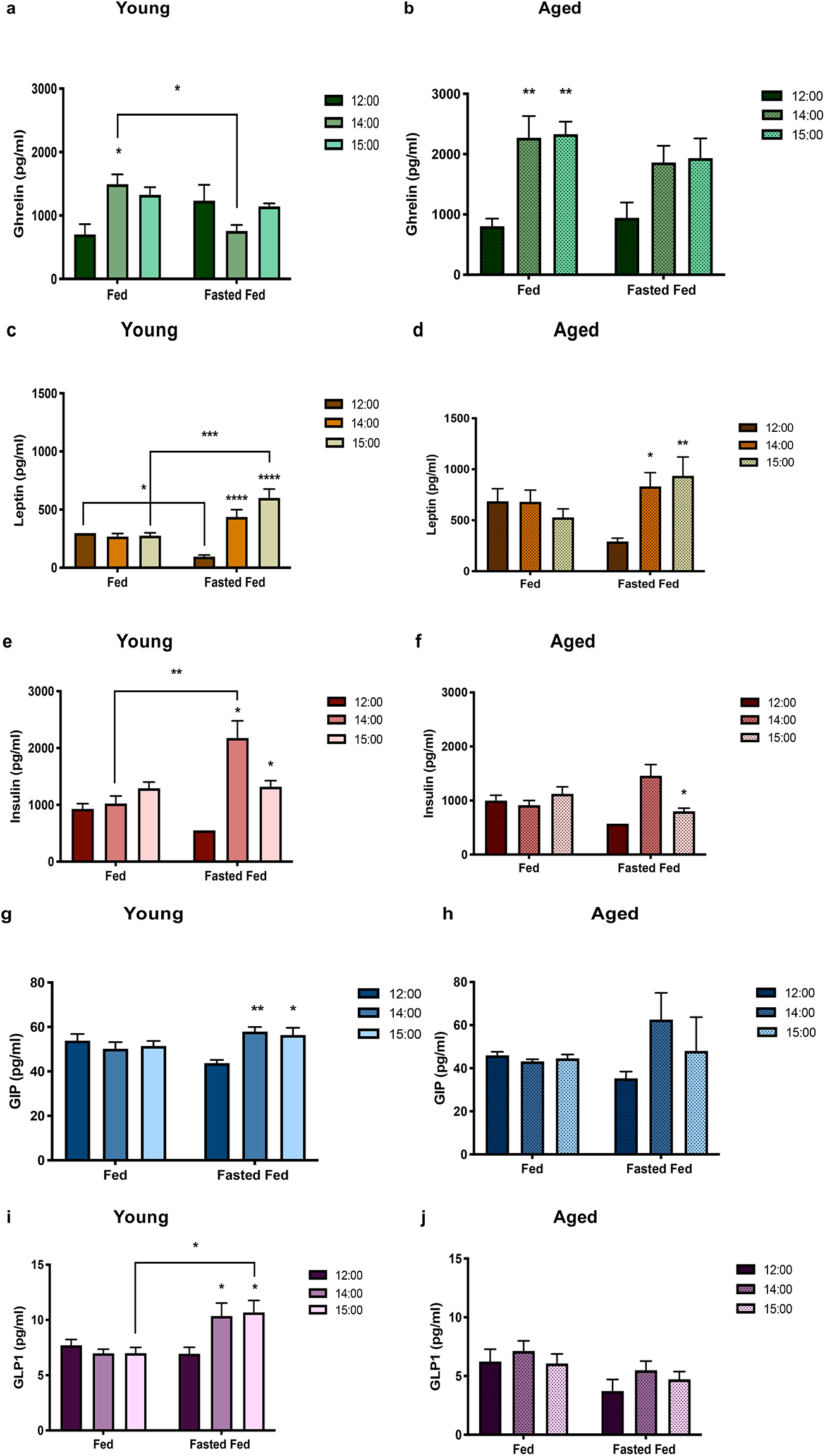
Serum concentrations of ghrelin, leptin, insulin, GIP and GLP1 in young adult 17-week-old and aged 20-month-old fed or overnight fasted 2-hour fed male C57BL/6 mice. **a**Serum ghrelin concentration significantly increased from 12:00 to 14:00 in young fed mice and declined at 14:00 in young fasted fed mice compared with that in fed mice. **b** Serum ghrelin concentrations were significantly higher at 14:00 and 15:00 compared to those at 12:00 in aged fed but not fasted fed mice. **c, d** In young and aged fasted fed mice, serum leptin concentrations significantly increased at 14:00 and 15:00 compared with fasting serum concentration at 12:00 and remained relatively unchanged in ad-lib fed mice. Concentrations of serum leptin was significantly lower at 12:00 and significantly higher at 15:00 in young fasted fed mice compared to those in ad-lib fed mice (c). **e** Serum insulin concentrations significantly increased from 12:00 to 14:00 in young fasted fed mice and in comparison to that in ad-lib fed mice, then significantly decreased at 15:00. **f** In aged fasted fed mice, serum insulin concentrations increased but not significantly from 12:00 to 14:00, then decreased significantly at 15:00 compared with that at 14:00 and remained unchanged in fed mice. **g, i** In young fasted fed mice, serum GIP and GLP1 concentrations significantly increased at 14:00 and 15:00 compared with fasting concentrations at 12:00 and remained unchanged in fed mice. Serum GLP1 concentrations were significantly higher at 15:00 in fasted fed mice compared to that in ad-lib fed mice (i). **h, j** There were no significant changes in serum GIP and GLIP1 concentrations within and between aged fed and fasted fed mice. All data are means ± SEM. *n* = 6 ad-libitum fed mice, *n* = 7 overnight (16-hour) fasted 2-hour fed mice. Two-way ANOVA with post-hoc Tukey comparisons; * *p* < 0.05, ** *p* < 0.01, *** *p* < 0.001, **** *p* < 0.0001

## Discussion

These data provide compelling evidence for a link between variations in timing of feeding and adaptive responses of bone to loading. Age-related bone loss across multiple sites within the skeleton suggests that systemic regulation plays an essential role in bone homeostasis. Concentrations of a number of gastro-entero-pancreatic hormones change before, during and after feeding, which have direct or indirect effects on bone cells. We interpret the data from this study to provide proof-of-principle that the effects of anabolic hormones in increasing bone formation in response to mechanical loading may be manipulated by changing timing of feeding in relation to exercise imposition. It has been previously shown that parathyroid hormone (an osteotropic hormone) administration potentiates effects of loading(14–16), but this study is the first report of a non-pharmacological intervention with such effects. The ability of feeding interventions to amplify the effects of loading stimuli is compelling, but its ability to turn a sub-threshold stimulus (one which alone does not stimulate any adaptive bone responses) into one causes significant new bone formation has potentially pervasive implications.

Our study of the fasting/2-hour feeding before mechanical loading in older mice did not recapitulate results seen in younger animals. Measurement of hormones in our studies gives clues to possible explanations for differences in adaptive responses of bones between young and aged mice as in the data on 5 different hormones, we determined clear differences in changes over the fasting/feeding/loading period in the two age groups. Age-related impairment of the function of mechanostat and decline of the skeletal robustness may have also reduced sensitivity of the cortical bone to mechanical stimulation in aged male mice. Further, the altered cortical bone response to the strain-related stimulus in aged male mice compared to young ones may reflect the deficiencies in osteoblast recruitment, differentiation and function. However, one limitation of this study is that we did not examine different timings of fasting and feeding in old mice as we had in young mice.

We were conscious that repeated blood sampling from mice could be stressful, and that such stress might affect levels of the hormones we wished to measure. However, assessments of serum cortisol concentrations during those blood sampling studies showed no significant changes suggesting minimal impact of the procedure on stresses experienced by the mice (Supplementary Figure S2).

Currently, although the effects we demonstrate are potent and robust, we have only association between the changes in bone responses and the hormonal alterations before, during and after feeding. There are however, compelling data on the way that such hormonal changes occur, and it is appropriate to consider some of the possible candidates in this connection.

Leptin is well recognised as a bone anabolic hormone, with its own effects mediated by central and peripheral mechanisms as well as those through other central relays^26^. We found like others before, that leptin concentrations are lower in fasted mice and that they increase following feeding^27–29^. It is known that leptin increases bone formation^30,31^, although to our knowledge, there are no reports of concurrent administration of leptin and loading of bones.

Ghrelin is thought to be antagonistic to leptin, has direct effects on bone homeostasis both *in vivo* and *in vitro*^32,33^ and changes during fasting and in advance of feeding^34,35^. The high circulating ghrelin concentrations during fasting and then low concentrations after food intake would be consistent with an influence of the leptin/ghrelin axis on the loading related bone formation that we demonstrate.

Insulin increases bone formation and decreases bone resorption both *in vivo* and *in vitro*^36–38^. During fasting circulating insulin concentrations are low (basal condition) and as eating commences, plasma insulin concentrations rise to regulate blood glucose concentrations. These elevated levels would be consistent with a potentiated anabolic bone response to loading. Specifically, we demonstrate that 2-hours of feeding after fasting increased serum insulin concentrations in young mice. The lack of similar change in insulin in the older mice is consistent with our demonstration that the same potentiation did not occur in them. As insulin has promiscuous effects on IGF receptors, which are expressed on osteoblasts^39^, it is likely that any insulin-mediated effect is complex.

GIP is a bone anabolic *in vitro*^40,41^ and mice overexpressing GIP have increased bone mass^42^. Similarly, GLP1 receptors are expressed on osteoblasts, bone marrow stromal cells^43^ and on thyroid C cells^44^, so GLP1 has the potential for direct and indirect actions on bone. However, as the concentrations of both hormones were unchanged in aged mice by our interventions we conclude that impairment of GIP or GLP1 signalling in bone cells are likely to be involved in attenuated bone response we observed. The roles of these different hormones could be clarified by interventional studies where hormones were administered in order to mimic the fasting/feeding changes, or in knockout or over-expressing mice, but those studies are beyond the scope of this research and any single report.

While fasting mice for 16 hours is a much more significant period of food deprivation than overnight fasting in humans, it is possible that similar biological interactions occur in humans, that could be exploited to benefit bone health. Future studies may allow us to determine whether such potentiating effects are observed in humans and what recommendations can be made to humans with respect to timing, amount and composition of food ingestion to obtain the maximal benefits from exercise. If we can identify interventions that potentiate bone’s response to exercise in young humans, then such interventions would increase adult bone mass so that with ageing, the additional strength of bones would extend time before osteoporotic “fracture thresholds” were reached. If similar interventions were identified in older mice and then humans too, then they could be translatable into a non-pharmacological way to improve bone health of older people. There are already widespread fasting and caloric restriction interventions in use in humans for weight control and for healthy ageing^45^. In particular, the 5:2 diet in which food intake is restricted to 500 – 600 calories on two days per week, which could be exploited to provide the equivalent of fast experienced by mice in this study. The likelihood that effects we show here could be translatable to humans has been increased by a recent study^46^ that showed in postmenopausal women eating only twice daily, positive effects of treadmill exercise on serum biomarkers of bone formation, one hour after eating but not 1 hour before eating.

## Abbreviations

Ct.Th: cortical bone thickness
BFR: bone formation rate
GIP: lucose-dependent insulinotropic polypeptide
GLP1: glucagon-like peptide-1
GLP2: glucagon-like peptide-2
MAR: mineral apposition rate
MS: mineralising surface
Tb.Pf: trabecular bone pattern factor

## Acknowledgements

This study was supported by CIMA (The MRC – Versus Arthritis Centre for Integrated Research into Musculoskeletal Ageing) (to TMS and JCM; MR/K006312/1) performed as a collaborative project between the Universities of Sheffield, Newcastle and Liverpool. The authors thank the members of the SkeletAL laboratory, The University of Sheffield, for tissue processing, Dr Carmen Martin-Ruiz (NIHR Newcastle Biomedical Research Centre, Ageing Research laboratories) for the analysis of serum cortisol concentrations in young adult and aged mice.

## Author Contributions

H. J. Samvelyan performed the research. T. M. Skerry conceived the original idea for the study, H. J. Samvelyan, J. C. Mathers and T. M. Skerry all contributed to the research design, analysis and interpretation of the data, drafted, reviewed, edited and approved the version of the manuscript for publication.

