## Supplemental data for "Feeding intervention potentiates the effect of mechanical loading to induce new bone formation in mice"

**<u>Supplemental material</u>**

**Table S1 Nutrient data of Teklad Global 18% Protein rodent diet (Teklad Diets, Madison, WI, USA) consumed by young adult and aged male C57BL/6 mice**

**Table S1**

Standard product form: Pellet

| **Macronutrients** | | |
|---|---|---|
| Crude Protein | % | 18.6 |
| Fat (ether extract) | % | 6.2 |
| Carbohydrate (available) [a] | % | 44.2 |
| Crude Fiber | % | 3.5 |
| Neutral Detergent Fiber [b] | % | 14.7 |
| Ash | % | 5.3 |
| Energy Density [c] | kcal/g (kJ/g) | 3.1 (13.0) |
| Calories from Protein | % | 24 |
| Calories from Fat | % | 18 |
| Calories from Carbohydrate | % | 58 |

[a]Carbohydrate (available) is calculated by subtracting neutral detergent fiber from total carbohydrates

[b] Neutral detergent fiber is an estimate of insoluble fiber, including cellulose, hemicellulose and lignin.

[c] Energy density is a calculated estimate of *metabolisable energy* based on the Atwater factors assigning 4 kcal/g to proteins, 9 kcal/g to fats and 4 kcal/g to available carbohydrates.

**Figure S1 Experimental design of mechanical loading, fluorescent labeling schedule and culling of young and aged male C57BL/6 mice**

Groups of young or aged male C57BL/6 mice were either fasted overnight for 16 hours or allowed free access to food ad libitum. At the end of fasting, mice were given access to food for 2 hours. The right tibia of each mouse was subjected to dynamic axial mechanical loading on alternate days for 2 weeks under the isoflurane-induced anaesthesia. The left tibiae were non-loaded controls in each animal. Calcein was injected subcutaneously into the mice on day 1 and day 12 of loading. All mice were culled on day 15. Experiment 1: 17-week-old mice loaded at 13N to induce peak strain magnitude of 2200 microstrain; experiment 2: 17-week-old mice loaded at 11N to induce peak strain magnitude of 1300 microstrain; experiment 3: 17-week-old mice loaded at 8N to induce peak strain magnitude of 1100 microstrain; experiment 4: 20-month-old mice loaded at 10N to induce peak strain magnitude of 2200 microstrain. Group sizes were $n = 7$ for experiments 1, 2 and 3, and $n = 8$ for experiment 4 respectively.

**Figure S2 Serum cortisol concentrations at three time points in young adult and aged male C57BL/6 mice**

Blood samples were obtained from the lateral tail vein of young adult 17-week-old ($n = 5$) and aged 20-month-old mice ($n = 7$) by tail incision method at 12:00, 14:00 and 15:00 of the experimental day. The concentrations of cortisol were measured in mouse sera using solid phase competitive ELISA assay. There were no significant changes in cortisol dynamics throughout the experiment in young adult and aged mice. Samples (ng/ml) were assessed in duplicate. All data are means ± SEM.

**Figure S1**

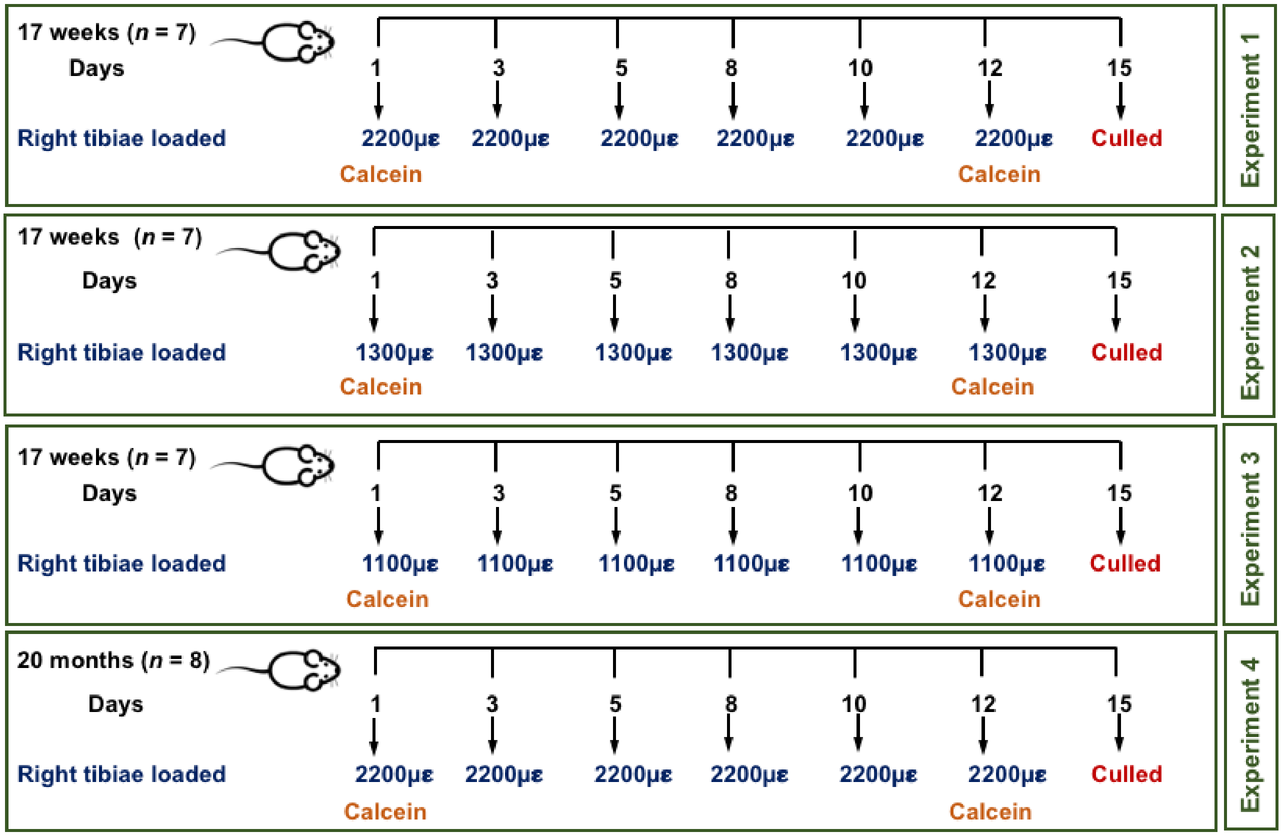


**Figure S2**

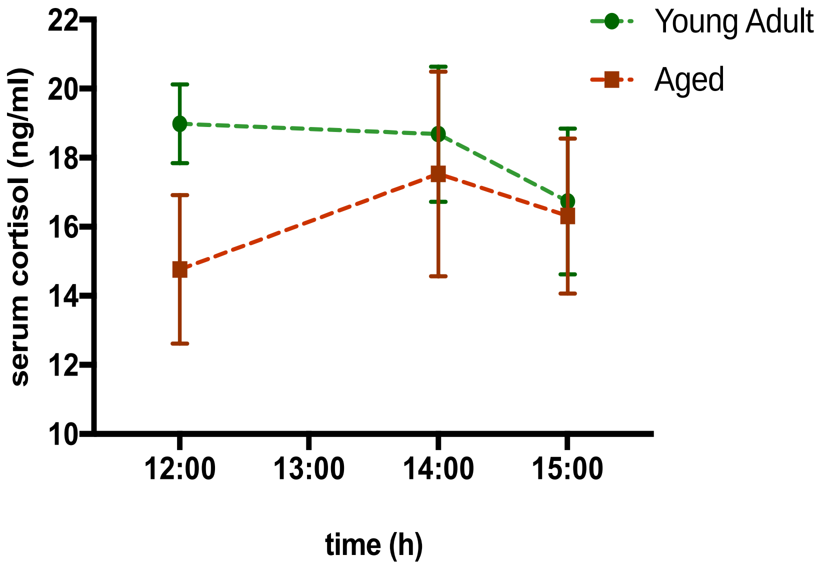
